# Deciphering rapid cell signaling and control of cell motility by reverse opto-chemical engineering

**DOI:** 10.1101/2024.02.26.582067

**Authors:** H. Hamzeh, A. Gong, M. Balbach, D. Fridman, H.G. Körschen, R. Pascal, F. Lavryk, A. Rennhack, R. Seifert, A. Hernandez-Clavijo, S. Pifferi, V. Dusend, B.K. Fleischmann, P. Sasse, A. Menini, B.M. Friedrich, L. Alvarez, U.B. Kaupp

## Abstract

Cells transform complex environmental stimuli into physiological responses. For time-varying stimuli or motile cells, the perception of the environment depends on the temporal stimulus pattern and cell motion, respectively. Here we report a concept, “reverse optochemical engineering” (ROCE), that uses temporal light patterns and photo-triggers to expose cells to virtual sensory landscapes while recording in real time their physiological responses and motor behavior. We studied cyclic-nucleotide signaling in cell lines, sperm, olfactory neurons, and cardiomyocytes. The technique can be employed for remote control of motility by light. We reprogrammed sperm from a chemotactic to a ‘phototactic’ cell that is attracted towards light. The method provides new opportunities to decipher the mechanisms and signaling molecules underlying rapid cellular computations, and thus reveal the wire diagram of cellular networks.

## Introduction

Biological cells are exposed over multiple timescales to numerous chemical and physical cues that ultimately evoke a cellular response or control behavior. A central aim of biology is to understand the transfer function between stimulus input *s*(*t*) and response output *o*(*t*) in quantitative and molecular terms. This endeavor is challenging because cells encounter a rich, spatially complex, and highly dynamic chemical environment composed of puffs, plumes, ramps, periodic cycles, turbulences, or voids of diverse stimuli (Sengupta et al., 2017; Taylor and Stocker, 2012). The complexity is further enhanced by feedback mechanisms that change the stimulus *s*(*t*). For example, stimulation of specific cells by a ligand may initiate down-regulation of ligand synthesis by other distant cell types. Another type of feedback results from cell motility. Motile cells probe a spatial distribution of sensory cues c(**r**) along their path **r**(*t*). Movement translates the spatial information into a temporal stimulus *s*(*t*), and cells use this sensory percept to continuously update their path accordingly (Fig. 1a,b). Such feedback by which a chosen motor action determines the future sensory input is called ‘information self-structuring’ (Lungarella and Sporns, 2006). The ultimate goal is to map in real time the biochemical circuits underlying navigation and the ensuing behavioral feedback while cells move in a sensory landscape. Yet, emulating a naturalistic sensory landscape in the laboratory is experimentally challenging.

**Figure 1:**
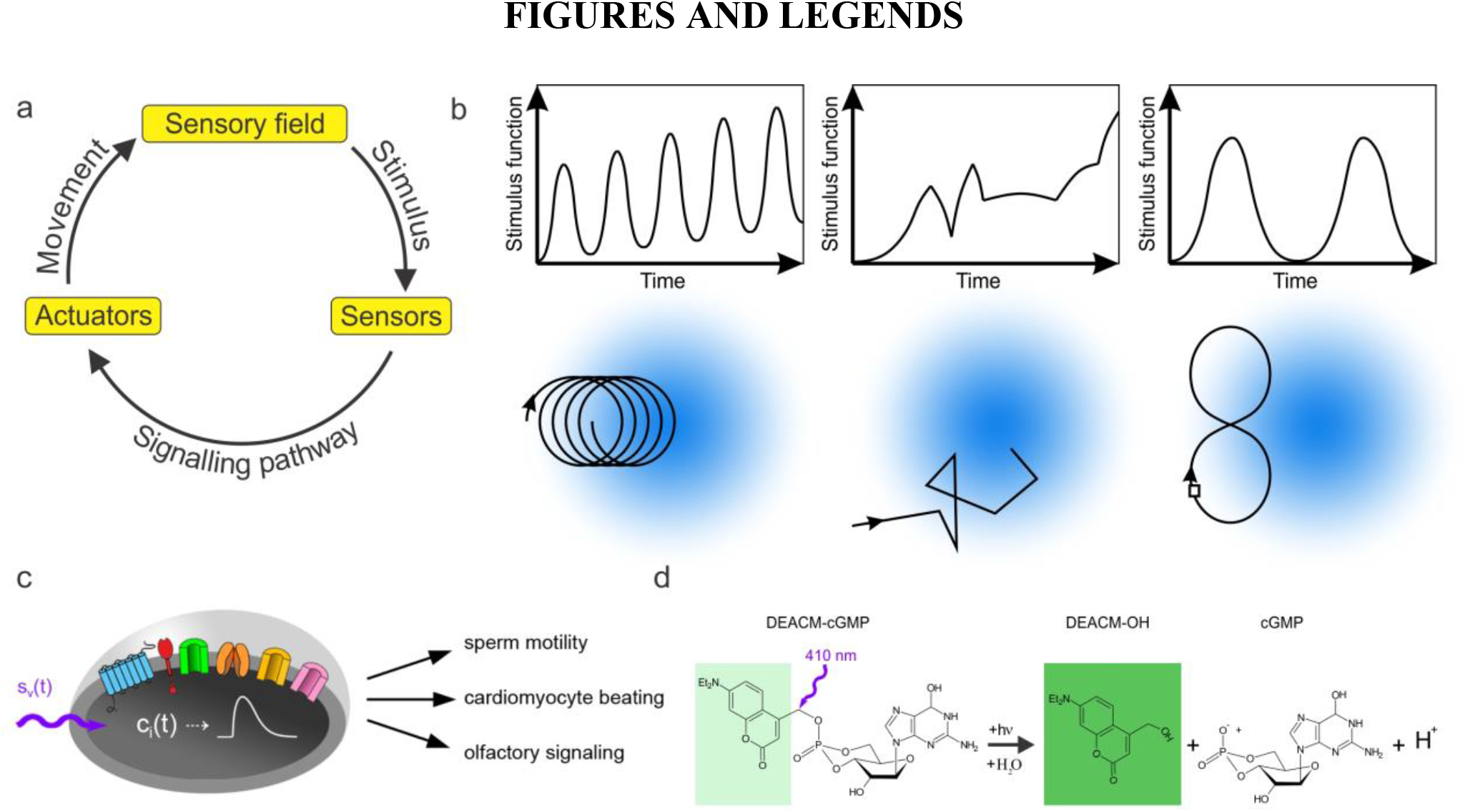
Recursive interactions between a sensory landscape, cellular signalling, and the motion path. **a**, The sensorimotor loop. Cells capture information from their environment. A signalling pathway translates the stimulus registered by the cell sensors into a motor output. As the cell moves in a sensory field, the stimulus function *s*(*t*) is also altered. Thus, *s*(*t*) is not only determined by the sensory field, but also by the active motion of the cell. **b**, Examples of temporal stimuli (top) that result from three different paths inside the same stimulus field (bottom). The stimulus field is shown in blue shades. **c**, Opto-chemical engineering of cellular messengers *c*_i_*(t)* (violet arrow) bypasses membrane receptors and imposes a virtual stimulus *s*_v_(*t*) on the cell. Icons represent cellular signaling components that control *c*_i_*(t)*. **d**, Chemical structure of DEACM-caged cGMP. The caging group (light green) has low background fluorescence. A 410-nm light flash photo-chemically cleaves the covalent bond between the cage and cGMP. The fluorescence increase of the free DEACM-OH cage (dark green) allows for quantification of the release.

A case in point is the directed movement of cells in a chemical gradient – called chemotaxis. Many organisms and cells are attracted by chemical factors that signal food, mates, or a favorable environment in general. Moving on periodic paths - drifting circles, helices, or zigzag movements - evolved as an efficient navigation strategy shared by cells and animals to locate a target (Brokaw, 1958; Clement et al., 2023; Crenshaw, 1996; Fenchel and Blackburn, 1999; Gomez-Marin et al., 2011; Huston et al., 2015; McHenry and Strother, 2003; Thar and Fenchel, 2001). Periodic locomotion patterns are also observed for other sensory taxes such as phototaxis (Jekely et al., 2008; Polin et al., 2009) and during active exploratory whisking of rodents and marsupials (Mitchinson et al., 2011). The stimulus functions *s*(*t*) generated by periodic movements are also periodic with typical frequencies ≥ 1 Hz.

A powerful approach for probing cellular signaling networks is to combine time-varying perturbations with signaling read-outs from live cells (Mettetal et al., 2008; Muzzey et al., 2009; Rahi et al., 2017; Shimizu et al., 2010; Toettcher et al., 2011). This strategy was previously used to study slow cellular processes, like gene expression or filopodia outgrowth that proceed on a minute-to-hours timescale and, therefore, are experimentally accessible by intermittent stimulation and time-lapse recordings. However, many biochemical signaling events related to sensing occur in less than a second.

Here, we extend this successful strategy to the millisecond time range using various waveforms of light and photo-triggers of the ubiquitous cellular messengers cAMP and cGMP (Fig. 1d). This method allows creating a *virtual* stimulus input *s*_v_(*t*) (Fig. 1c) that mimics the stimulus encountered in natural sensory landscapes. The concept is reminiscent of reverse genetics, which introduces specific mutations into genes and analyses the resulting phenotype; we term this approach accordingly reverse opto-chemical engineering (ROCE). While stimulating cells with temporal light patterns, we simultaneously recorded cellular responses using orthogonal potentiometric probes and Ca^2+^ indicator dyes. We study signaling events in cell lines, cardiomyocytes, olfactory neurons, and sperm. This approach revealed the sensorimotor transfer function of sperm chemotaxis. We used ROCE to control sperm motility by light and to ‘photoprint’ spatial patterns of sperm cells.

## Results

### Implementation of reversed opto-chemical engineering

A virtual stimulus function *s*_v_(*t*) could be established either by caged chemical compounds or optogenetic tools. We opted in favor of caged compounds and against optogenetic tools for the following reasons.

Genetically encoded light-sensitive enzymes for manipulation of cyclic-nucleotide levels are available, notably photosensitive adenylate cyclase bPAC (Jansen et al., 2015; Schröder-Lang et al., 2007; Stierl et al., 2011), rhodopsin/guanylate cyclase fusion proteins RhGC (Avelar et al., 2014; Gao et al., 2015; Scheib et al., 2015), and light-activated phosphodiesterases (LAPD) (Gasser et al., 2014; Scheib et al., 2015). However, for several reasons these enzymes are not ideal for mimicking and rigorous quantification of sub-second signaling events: copy numbers and folding of proteins vary and are difficult to determine; the relationship between light energy and activity is complex or variable; the activation/deactivation kinetics of the respective enzymes is too slow for probing millisecond signaling events (Jansen et al., 2015; Stierl et al., 2011); and some light-activated enzymes, like bPAC and LAPD, exhibit basal activity that interferes with the resting state of cells. Instead, we chose DEACM-caged cNMPs to rapidly and quantitatively manipulate cAMP and cGMP levels by light (Hagen et al., 2002) (Fig. 1d). DEACM-caged cNMPs are completely inactive; their high quantum yield (0.27) and activation spectrum (λ_peak_ = 410 nm) (Hagen et al., 2005) allows using low, non-toxic light levels. Owing to their ultrafast photochemistry (≤ 10 ns) (Eckhardt et al., 2002), physiological reactions can be triggered with sub-millisecond time resolution. Moreover, our caged NMPs offer excellent spectral properties: they do not require UV light for release and allow parallel monitoring of multiple cellular reactions using fluorescent dyes or proteins without release, i.e., stimulating and measuring the cellular response can be achieved independently (Kierzek et al., 2021). This cannot be achieved with optogenetic tools, such as channel rhodopsin, as their action or response spectra overlap with the excitation spectrum of most fluorescent reporters (Lin, 2011). Finally, photo-release of cNMP can be quantified and calibrated (see Methods).

We tested ROCE using three prototypical waveforms *s*_v_(*t*): (1) a square pulse, (2) a sinusoidal waveform, and (3) a linear ramp (Fig. 2a). For a stationary cell, the waveforms mimic a step increase of stimulus, a periodic or pulsatile stimulus, and a steadily increasing stimulus, respectively. The cumulative light-stimulated cAMP increase should follow the integral

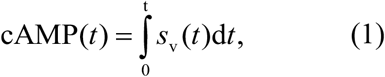

where the virtual stimulus *s*_v_(*t*) = *k*_rel_ *I*(*t*) is the product of the rate constant *k*_rel_ of photo-release and the intensity *I*(*t*) of the photolyzing light. The fractional photo-release of cAMP was monitored *in vitro* using the enhanced fluorescence of the free DEACM-OH cage (Eckhardt et al., 2002) (Fig. 2a) and in CHO cells using the cAMP sensor cADDis (see Methods) (Fig. 2b). The prediction embodied in eqn. (1) is borne out by experiment *in vitro* and *in vivo*: the three waveforms produced a linear, a frequency-modulated linear, and a quadratic rise of cAMP concentration, respectively (Fig. 2a,b and Supplementary Fig. 1a). At longer stimulation times, the *in vivo* cADDis signal in CHO cells leveled off (Fig. 2b, middle panel) because the sensor becomes saturated with cAMP, as predicted by a simple binding model (see Methods). After the light stimulus has been switched off, the cADDis fluorescence remained elevated for many seconds, indicating that, on this time scale, CHO cells lack tangible PDE activity that hydrolyzes cAMP. Similar results were obtained by imaging cAMP levels in single cells (Fig. 2c and Supplementary Fig. 1b). In conclusion, the changes in cAMP observed by experiment and predicted by theory perfectly match. Next, we tested ROCE in three different cell systems that employ cyclic-nucleotide signaling to register temporal stimulus patterns: cardiomyocytes, olfactory sensory neurons (OSN), and sperm.

**Figure 2:**
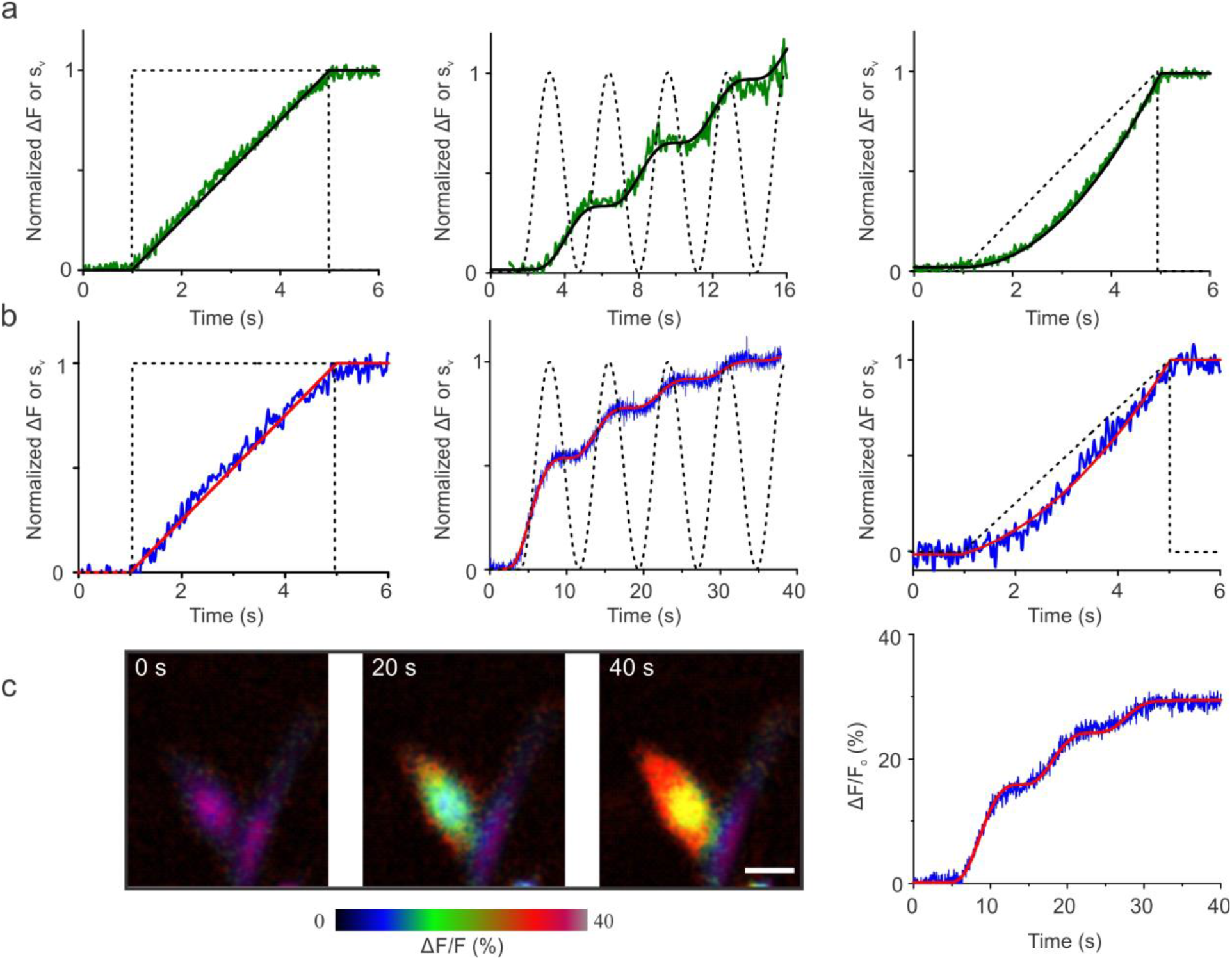
Implementation of the ROCE method. **a**, Time course of DEACM-OH fluorescence (green) during photolysis using a virtual stimulus *s*_v_(*t*). Waveforms of *s*_v_(*t*) are shown (dashed line). The calculated cumulative cAMP release for the various *s*_v_(*t*) waveforms is shown in black. Data for *in vitro* experiments with DEACM-caged cAMP (15 μM) in the buffer; λ_exc_ = 365 nm; λ_em_ = 490 nm. **b**, Fluorescence time course of the cAMP sensor cADDis (blue) evoked by *s*_v_(*t*) in caged cAMP-loaded CHO cells (15 μM); fit using a modified simple binding isotherm is shown in red (see Methods). **c**, Single-cell imaging. Left: false-color images of cADDis-expressing HEK cells showing the increase of fluorescence at three time points upon photo-release of cAMP by a periodic input *s*_v_(*t*). Scale bar 20 μm. Color scale bar indicating the cADDis fluorescence intensity is shown below the false-color images. Right: fluorescence time trace (blue) from a single cell. The red line is a fit of a modified simple binding isotherm.

### Application of ROCE to cardiomyocytes

Adrenergic signaling enhances spontaneous periodic activity of cardiac pacemaker cells by elevating cAMP levels. Direct binding of cAMP to hyperpolarization-activated, cyclic nucleotide-gated (HCN) pacemaker channels accelerates the heartbeat. Moreover, protein kinase A (PKA)-mediated activation of L-type Ca^2+^ channels, SERCA2a pump, and other proteins also accelerate the beat rate by voltage- and Ca^2+^-dependent clock mechanisms (Lakatta and DiFrancesco, 2009). HCN channels and PKA *in vitro* are activated by approximately 200-nM and 10-nM cAMP concentrations, respectively. However, the precise cAMP concentrations required to activate these targets in a living cell are not known. Here, we use precise quantification of intracellular cAMP release to measure the range of cAMP concentrations that affect the beat rate. Single cultured neonatal mouse cardiomyocytes were loaded with DEACM-caged cAMP (20 μM), and spontaneous beating frequency was recorded using infrared video microscopy (Bruegmann et al., 2010). Release of cAMP was controlled by precisely timed light pulses (λ_exc_ = 385 nm, 1 to 250 ms), corresponding to calculated cAMP concentrations from 0.32 to 20 μM. Approximately 50% of cardiomyocytes (38 of 74) responded with an increase of the beat frequency (Fig. 3a left). The beat frequency gradually increased with cAMP concentration, being half-maximal at a cAMP concentration (*K*_1/2_) of 3.57 ± 1.16 μM (*n* = 3 individual cell isolations) (Fig. 3a right). This value is approximately 17-fold larger than the *K*_1/2_ constant of HCN activation by cAMP measured in excised patches of cardiac pacemaker cells (0.21 μM) (DiFrancesco and Tortora, 1991) and about 350-fold larger than the *K*_1/2_ of PKA activation, suggesting that cAMP hydrolysis by phosphodiesterase activity in intact cardiomyocytes plays an important role for the adrenergic cAMP response (Koschinski and Zaccolo, 2017). Whereas ROCE is relatively easy to establish in single cells, cell layers, or cell suspensions, preliminary experiments suggest that ROCE also holds promise in Langendorff preparations of the beating heart. All-optical electrophysiology (optogenetics) has been also applied to the zebrafish beating heart (Jia et al., 2023).

**Figure 3.**
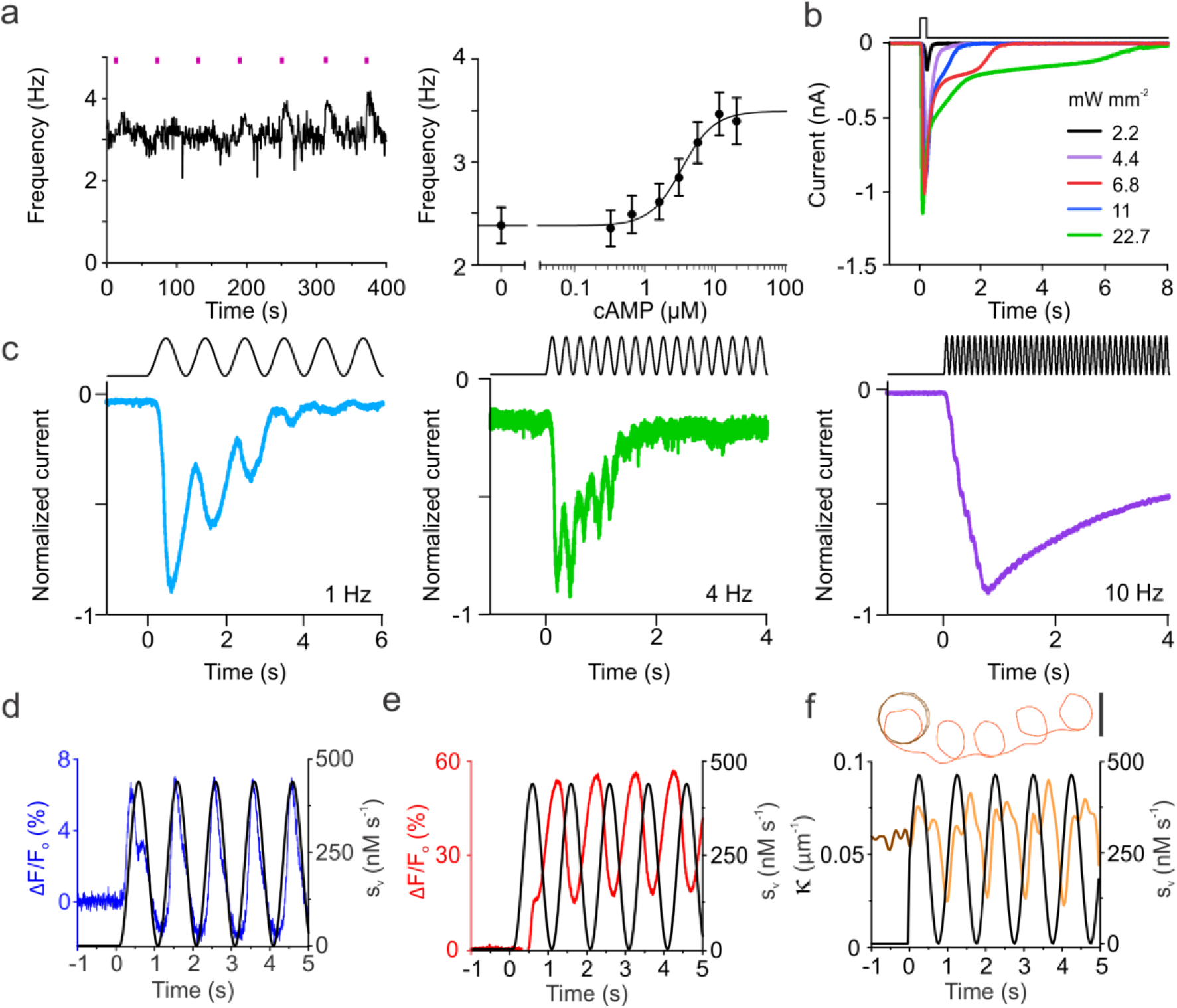
Applying ROCE to cyclic-nucleotide signaling in cardiomyocytes, olfactory sensory neurons, and sperm. **a**, Modulation of beating frequency in isolated neonatal mouse cardiomyocytes upon release of cAMP. Pulse durations ranging from 1 to 250 ms were used, equivalent to 0.32 up to 20 μM of cAMP (left). Dose-response curve of the change in beating frequency by cAMP (right). Data was fitted to a logistic function with *K*_1/2_ = 3.57 ± 1.16 μM (*n* = 3 cell isolations, a total of 37 cells was analyzed). **b**, Current responses induced in olfactory sensory neurons by cAMP uncaging with light pulses (top) of the indicated power density. Caged cAMP (20 μM) diffused into the cell from the patch pipette, and flashes of 200 ms duration were applied to the ciliary region. Whole-cell current responses were measured at the holding potential of −80 mV. The upper trace indicates the time of light application. **c**, Normalized current responses in different OSNs induced by a sinusoidal light stimulus (top, black) at the indicated frequencies of 1, 4, or 10 Hz. The peak power density was 2.2 mW mm^-2^. The maximal amplitude varied between 80 pA to 600 pA. **d**, Changes in voltage V_m_ (blue) of sperm evoked by a 1-Hz sinusoidal light stimulus *s*_v_(*t*) (black); responses were registered with the fluorescent potentiometric probe FluoVolt. **e**, Changes in [Ca^2+^]_i_ (red) registered with the Ca^2+^ dye Fluo-4. The light power for photolysis was 0.24 mJ, equivalent to the release of 220 nM cGMP s^-1^. **f**, Changes in 2D path curvature κ(*t*) upon stimulation by a 1-Hz sinusoidal *s*_v_(*t*); κ(*t*) was derived from the path of sperm swimming at a glass/water interface (upper); basal circular path over time (brown), swimming path after onset of stimulation (orange). Scale bar 50 μm.

### Application of ROCE to olfactory neurons

Next, we examined cAMP signaling in OSNs. A cAMP-signaling cascade transduces the binding of odorant molecules to receptors into an electrical response. Signaling occurs in specialized cilia, where the increase of cAMP activates cyclic nucleotide-gated (CNG) channels allowing Ca^2+^ influx. The increase of intra-cilia Ca^2+^ concentration ([Ca^2+^]_i_) activates a Ca^2+^-dependent chloride current that further amplifies the electrical signal (Dibattista et al., 2017). Using the whole-cell patch-clamp technique, light-evoked currents were recorded from OSNs loaded with DEACM-caged cAMP. Brief light pulses of 200-ms duration applied to the ciliary region of neurons elicited a fast deactivating current that was dose-dependent and saturated at 6.8 mW mm^-2^ of light power density (Fig. 3b). A further increase of light energy produced the same peak current but prolonged the time necessary to return to baseline levels. The light-induced currents showed the typical OSN response to a fast increase of [cAMP]_i_ (Pietra et al., 2016).

Temporal fluctuations in odor concentration is used by organisms to gather information on odor plumes and their source (Wachowiak, 2011). Moreover, odorants are delivered to the olfactory epithelium during air inspiration and by sniffing. The respiratory frequency of mice is about 2 Hz at rest and can increase up to 10 Hz during active olfactory exploration (Ghatpande and Reisert, 2011; Wachowiak, 2011). We used ROCE to mimic stimulation due to periodic air inspiration. When OSNs were stimulated with sinusoidal 1-Hz light waveforms, the transduction currents oscillated almost in phase with the oscillatory stimulus (Fig. 3c, left panel). For higher stimulation frequencies of 4 and 10 Hz, the frequency-modulated response amplitude was attenuated (Fig. 3c, middle and right panel). Our results are consistent with data obtained using individual odorant pulses applied to isolated OSNs (Ghatpande and Reisert, 2011). The filter characteristics of OSNs could be shaped at several stages along the signaling pathway, notably by termination of odorant receptor activity, cAMP hydrolysis by PDE, or shutdown of CNG and Cl^-^ channels. Because the virtual stimulus bypasses odorant receptors and cAMP synthesis, our results suggest that the frequency dependence of odorant signaling is determined by the rate of recovery events downstream of the receptor and the adenylate cyclase (Ghatpande and Reisert, 2011). However, several obvious candidates like cAMP hydrolysis in the olfactory cilium by PDE1C (Cygnar and Zhao, 2009) or CNG channel shutdown (Song et al., 2008) contribute little if at all to response termination (Reisert and Zhao, 2011). Most likely, Ca^2+^ clearance by the NCKX4 exchanger controls recovery kinetics (Stephan et al., 2011). Future work using ROCE can test this hypothesis.

### Cellular and motility responses of sperm to periodic stimulation

Next, we tested ROCE to study and manipulate chemotactic signaling and swimming behavior of sea urchin sperm. Sperm cells swimming close to a boundary surface trace chemoattractant gradients along drifting circles (Alvarez et al., 2012; Jikeli et al., 2015; Kaupp et al., 2003), which leads to self-entrained stimulus oscillations with the frequency of circular swimming (approximately 1 Hz). A cGMP-signaling pathway (Supplementary Fig. 3) translates the periodic stimulation into bursts of [Ca^2+^]_i_ that modulate the flagellar beat (Alvarez et al., 2012). Three key signaling events – a rise of cGMP, a transient hyperpolarization ΔV_m_ of the cell membrane, and a rise of [Ca^2+^]_i_– all occur within 10 ms to 1 s (Hamzeh et al., 2019). To emulate *s*(*t*) experienced by a cell that moves along drifting circles in a chemical gradient, we stimulated caged cGMP-loaded sperm with sinusoidal waveforms of light and recorded changes in V_m_ and [Ca^2+^]_i_ using fluorescent dyes.

The sinusoidal *s*_v_(*t*) evoked oscillations of V_m_(*t*). Waveforms of stimulus and normalized response almost perfectly superimpose (Fig. 3d). The frequency-modulated amplitude of V_m_(*t*) was not attenuated during stimulation. Thus, at least for physiological 1-Hz stimulation used here, the voltage response does not adapt to oscillatory *s*_v_(*t*). The waveform of the Ca^2+^ response also recapitulated *s*_v_(*t*), but was phase-shifted by φ = 250º ± 13º (*n* = 4) (Fig. 3e). The phase shift results from the characteristic latency of the Ca^2+^ response (Kaupp et al., 2003; Seifert et al., 2015). The regular sinusoidal waveforms of V_m_(*t*) and Ca^2+^(*t*) responses suggest that sperm sample a stimulus continuously on a millisecond time scale rather than discretely at regular intervals. Thus, sperm transduce a time-continuous analog input signal *s*_v_(*t*) into a time-continuous analog output response. Finally, V_m_(*t*) and Ca^2+^(*t*) signals are phase-locked with the oscillating virtual stimulus *s*_v_(*t*).

We examined the swimming path produced by light-stimulated Ca^2+^ oscillations. A 1-Hz sinusoidal stimulus *s*_v_(*t*) was imposed onto caged cGMP-loaded sperm swimming in 2D at the glass/water interface of a shallow recording chamber (Fig. 3f; Supplementary Movie 1). Before stimulation, sperm swam along regular circles (Fig. 3f, bottom panel brown) with a constant path curvature κ (Fig. 3f, bottom panel brown). During sinusoidal light stimulation, sperm swam along drifting circles (Fig. 3f, top panel orange). The drift was previously explained as a result of oscillations of path curvature κ at the frequency of circular swimming (Fig. 3f, bottom panel orange (Alvarez et al., 2012; Friedrich and Jülicher, 2007). Thus, the natural chemotactic 2D navigation pattern in a concentration gradient is mimicked by periodic light stimulation, in the absence of any external chemical cues.

### Recording of transfer functions using ROCE

While a cell is moving, it translates a spatial concentration field *c*(**r**) into a temporal stimulus *s*(*t*). The acuity of a cell’s ‘spatial’ percept depends on the filtering characteristics of the signaling pathway that encodes the cellular response. We determined the transfer function **H** between the input *s*_v_(*t*) and the Ca^2+^(*t*) output of the signaling system using frequency sweeps ranging from *f* = 0.1 to 10 Hz (Fig. 4a). The amplitude of the Ca^2+^ response was maximal at *f* ≤ 1 Hz and dropped off at higher frequencies with a cut-off frequency *f*_c_ = 1.2 ± 0.16 Hz (*n* = 4) (Fig. 4b black). Thus, the signaling network serves as a low-pass filter with a cut-off just below the 1-Hz frequency of circular swimming (Alvarez et al., 2012; Jikeli et al., 2015). As a test, the transfer function **H** was validated using a stimulus *s*_v_(*t*) that represented the superposition of 1-Hz and 6-Hz signals. The Ca^2+^ signal and the response predicted by **H** matched (Fig. 4d). Similarly, we can define a second transfer function **H**_**κ**_ that links the stimulus and the motor response, i.e., changes in the path curvature κ of swimming sperm. We term this **H**_**κ**_ the stimulus-motor transfer function (Fig. 4b,c orange). Both transfer functions (**H**_**κ**_ for κ(*t*) and **H** for Ca^2+^(*t*)) displayed similar cut-off frequencies (Fig. 4b). Furthermore, the phase shift Δφ between Ca^2+^(*t*) and κ(*t*) was approximately 90° (Fig. 4c) as predicted by the transfer function between Ca^2+^(*t*) and κ(*t*) as previously reported (Alvarez et al., 2012). Finally, the phase shift Δφ between *s*_v_(*t*) and both Ca^2+^(*t*) and κ(*t*) decreased at higher frequencies.

**Figure 4.**
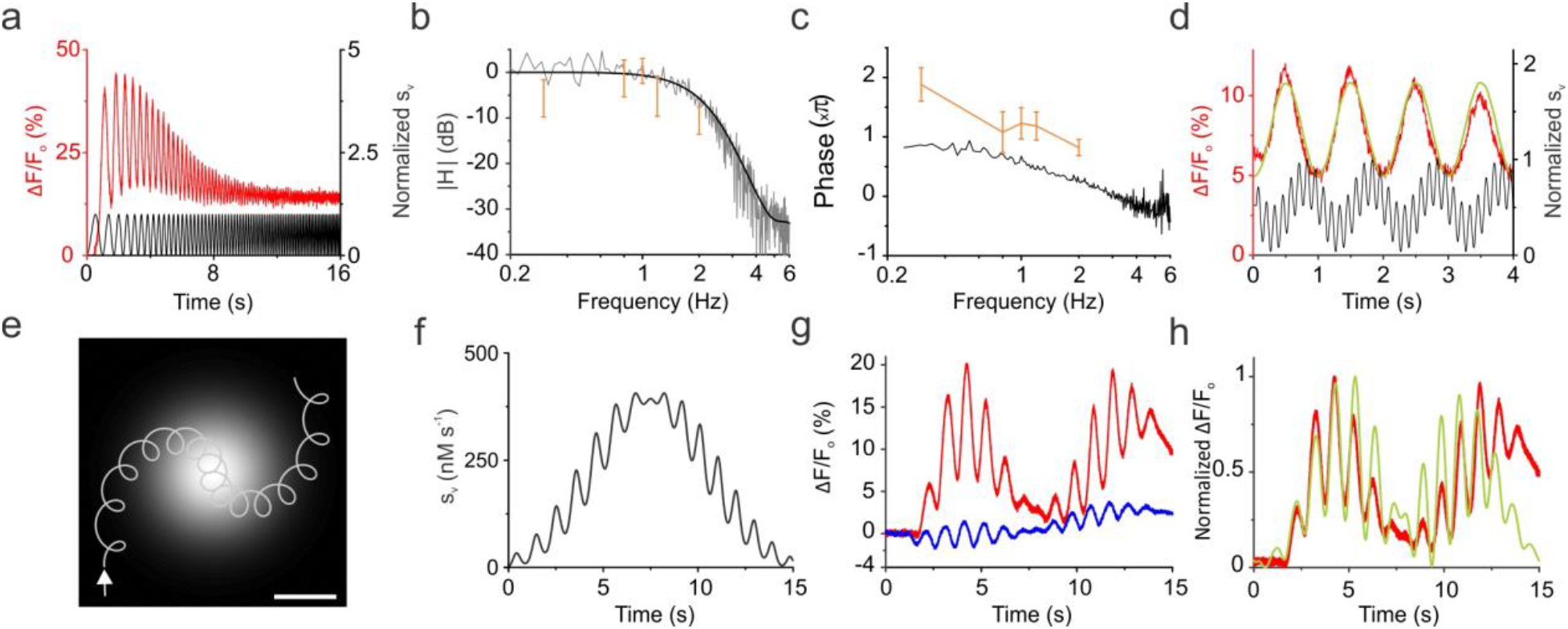
The transfer functions of chemotactic signaling. **a**, Frequency-dependent Ca^2+^ response (red) of sperm. Stimulus sweep *s*_v_(*t*) (black) used to determine the transfer function of the signaling pathway of sperm. ΔF/F_o_ indicates the changes in [Ca^2+^]_i_. **b**, Estimated magnitude of the raw (grey) and smoothed (black) transfer function |**H**(*f*)| of the Ca^2+^ response and of the curvature amplitude ± s.d. (orange). **c**, Phase relation of Ca^2+^ signal (black) and curvature (orange) as function of stimulus frequency. **d**, Validation of the transfer function using a test *s*_v_(*t*) that was the superposition of two sinusoidal oscillations of different frequencies (1 and 6 Hz) (black); for comparison, the Ca^2+^ signal (red) was superimposed on the output signal (light green) obtained by applying the transfer function to the input signal. **e**, Emulating chemotactic signaling in a virtual gradient. Hypothetical swimming path of sperm (light grey) while traversing a Gaussian-shaped concentration profile (grey scale); arrow shows the swimming direction (scale bar 100 μm). **f**, Virtual stimulus function *s*_v_(*t*) that emulates the stimulus while swimming in the Gaussian-shaped chemical field along the hypothetical path. **g**, Changes in voltage V_m_ (blue) and [Ca^2+^]_i_ (red) during navigation in a virtual gradient were recorded with FluoVolt and Fluo-4, respectively. **h**, Output Ca^2+^ signal (light green), predicted from the transfer function in part b and c, is superimposed on the measured Ca^2+^ signal (red). The predicted output is the result of convolution between *s*_v_(*t*) and the transfer function **H**.

Next, a virtual chemical landscape was emulated. We computed the stimulus that a sperm cell experiences as it navigates on drifting circles in a Gaussian profile of chemoattractant concentration, first moving up gradient and then down gradient (Fig. 4e,f). When the equivalent *s*_v_(*t*) was imposed onto sperm, V_m_ and Ca^2+^ oscillated with a characteristic waxing-and-waning pattern both during virtual swimming up- and down gradient (Fig. 4g). At the maximum of the light profile, corresponding to the peak of the virtual spatial profile, the amplitude of V_m_ and Ca^2+^ oscillations approached zero (Fig. 4g). The waxing-and-waning response pattern is also predicted by the transfer function **H** (Fig. 4h). Several insights are gained from the virtual-landscape experiment. Sperm keep their responsiveness to periodic stimulation over a wide range of mean stimulus strength. Moreover, phase-locking between the stimulus *s*_v_(*t*) and V_m_(*t*) and Ca^2+^(*t*) responses is preserved. Finally, sperm also display oscillatory V_m_ and Ca^2+^ responses when the mean stimulus strength decreases, which signifies a descent from high to low chemoattractant concentrations. Thus, sperm maintain their typical response required for positive chemotaxis, even for a virtual stimulus that corresponds to a hypothetical case.

### Optical control of cell motility

We sought further insight into stimulus-motor coupling during stimulation with physiological 1-Hz sinusoidal stimuli. For sperm swimming along circular paths, the stimulus traced from a concentration gradient oscillates precisely with the frequency *f*_0_ of circular swimming. Theory predicts that, during chemotaxis, sperm swim along drifting circles resulting from oscillations of the path curvature (Friedrich and Jülicher, 2007), and experiments confirmed this prediction (Alvarez et al., 2012; Böhmer et al., 2005)

ROCE allows breaking the natural self-entrainment between the stimulus frequency *f*_*s*_and the frequency *f*_0_ of circular swimming. The resultant swimming paths resemble ‘spirograph orbits’ (Fig. 5a, Supplementary Movie 1 and Supplementary Table 1) that are known from the children toy Spirograph, where a circle with a pen roles without slipping inside a larger circle. In mathematics, ‘spirograph orbits’ belong to a diverse class of plane curves, referred to as hypotrochoids (Fig. 5a). Extension of the previous theory predicts such ‘spirograph orbits’ if the stimulus frequency *f*_s_ does not match the frequency *f*_0_ of circular swimming (Fig. 5b, Supplementary Note 2). The calculated ‘spirograph orbits’ feature a different number of high-curvature ‘lobes’ that are located either inside (for *f*_s_ < *f*_0_) or outside (*f*_s_ > *f*_0_) the centerline of a circular path. The core idea of the theory is as follows. If the frequency *f*_s_ of the stimulus *s*_v_(*t*) and *f*_0_ of circular swimming perfectly match (*f*_s_ = *f*_0_), the cell velocity vector **v** points in the same direction before and after a full cycle of stimulus oscillations; consequently, swimming circles drift along a straight path (Fig. 5b and Supplementary Fig. 4). Thus, for self-entrained oscillations with *f*_s_ = *f*_0_, sperm move on drifting circles with an almost straight centerline. However, when *f*_s_ and *f*_0_ differ, **v** will adopt a non-zero net rotation after each stimulus period. Consequently, cells move on ‘spirograph orbits’, which may be considered as drifting circles with a circular centerline (Fig. 5b and Supplementary Fig. 4). The effective frequency of circular swimming for moving along this circular centerline equals the frequency difference |*f*_s_ – *f*_0_|. The theory also suggests that the collection of diverse paths (Supplementary Table 1) results from distributions of swimming velocities or circle diameter. The quantitative agreement between experimental and theoretical paths is excellent.

**Figure 5.**
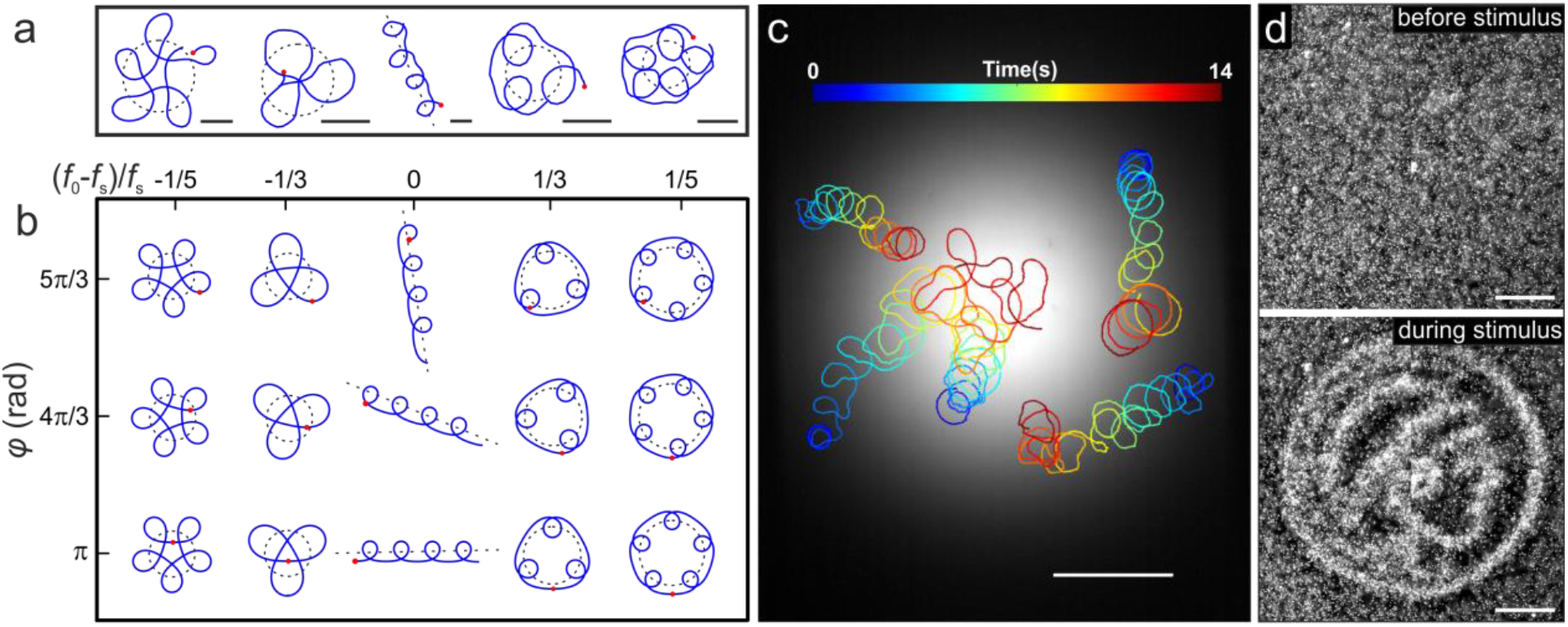
Control of sperm motility by light. **a**, Collection of experimental trajectories of caged cGMP-loaded sperm stimulated with 1-Hz sinusoidal UV light. Starting position: red dot; centerline of trajectories: black dashed lines. Scale bars: 50 μm. **b**, Predicted trajectories for cells swimming with a path curvature κ(*t*) that oscillates at the stimulation frequency *f*_s_ and with a phase φwith respect to the stimulus *s*_v_(*t*). Rational ratios between the frequency *f*_0_ of circular swimming and *f*_s_ determine the n-fold symmetry of trajectories (columns). Variations of φresult in identical patterns that are rotated (rows) (see Supplementary Note). **c**, Swimming paths of sperm swimming up a Gaussian-shaped light profile in the absence of chemoattractant. While swimming in the light profile, cGMP is released according to the local UV light power (grey shades); colored bar indicates the time; scale bar 200 μm. **d**, Remote control of sperm by light stimulation to draw a Minerva icon. Each white dot corresponds to a single sperm cell (imaged with dark-field microscopy). Pictures are shown before (top) and 5 s after switching on the UV light (bottom). Scale bars 500 μm.

ROCE enables directional control of motile cells by light. We imposed a spatial light profile on sperm swimming along circles (Fig. 5c). Under these conditions, the stimulus frequency is entrained to the circular-swimming frequency (*f*_s_ = *f*_0_), i.e., cell motion and stimulus become coupled. Using this ROCE paradigm combined with spatial light profiles, we ‘hijacked’ the chemotactic signaling network and reprogrammed sperm from a chemotactic to a ‘phototactic’ cell that is attracted towards light (Fig. 5c and Supplementary Movie 2). Using light patterns, it is possible to paint images by cell-density distributions (Fig. 5d and Supplementary Movie 3). Full remote control by various *s*_v_(*t*) designs will require monitoring the circular swimming frequency and adjusting the frequency of light stimulation in real time.

## Discussion

ROCE allows to expose motile cells to a time-dependent virtual stimulus *s*_v_(*t*) by using light and, at the same time, to record biochemical, electrical, and behavioral responses. Delivery of *s*_v_(*t*) is temporally precise, quantitative, and provides access to the millisecond time domain of physiological reactions. We demonstrate the potential of ROCE for the ubiquitous messengers cAMP and cGMP. Yet, ROCE is as versatile as the collection of available chemical photo-triggers, including caged versions of Ca^2+^, phosphoinositides, lipids, proteins, or CO_2_ (Goeldner and Givens, 2005). Optogenetic enzymes and sensors that operate on the millisecond time scale can further expand ROCE’s applicability. Moreover, several (orthogonal) photo-triggers could be used that can activate two cell components simultaneously but independently. Thereby, crosstalk between cellular network motifs could be studied.

ROCE can emulate complex natural sensory landscapes such as odor plumes or concentration fields distorted by turbulent flow that are otherwise difficult to establish and analyze quantitatively. ROCE can be employed to study cell responses evoked either by rapid cell movement in a stationary chemical gradient or, alternatively, by moving ‘wave fronts’ of chemical stimuli washing over a stationary cell. The washing-wave scenario is pertinent to delivery of metabolically active substances by the blood stream or fluid flow generated by ciliary beating in the olfactory epithelium or brain ventricles (Olstad et al., 2019; Reiten et al., 2017). Another important example is provided by slowly moving chemotactic cells exposed to moving wave fronts of chemoattractant, e.g. in the slime mold *Dictyostelium discoideum*. When a wave approaches, cells first experience an increase of chemoattractant concentration. Thereafter, as the peak of the wave front passes, cells are exposed to an oppositely directed gradient. Despite this reversal of the gradient, *Dictyostelium* cells continue to move in the same direction. How chemotactic cells can respond to a stimulus that increases in time, yet ignore a decreasing stimulus is known as the ‘back-of-the-wave’ problem (Huang and Iglesias, 2014; Skoge et al., 2014). One explanation assumes that cells adapt and become less sensitive while the stimulus increases. At the back of the wave, when the stimulus is decreasing again, desensitized cells cannot respond anymore. Alternatively, cells may develop an intrinsic polarity, a kind of molecular memory that allows motile cells maintaining their direction. By contrast, sperm do not polarize in a gradient, and the circular swimming of sperm in 2D does not possess persistence. In fact, sperm react to a concentration decrease almost instantaneously. Two fundamental gradient-sensing mechanisms can be distinguished: *spatial* and *temporal* comparison. Spatial comparison refers to the measurement of a spatial gradient across the cell diameter; temporal comparison refers to the detection of temporal changes of a stimulus while the cell actively moves inside a stimulus landscape. It can be rather difficult to distinguish spatial from temporal comparison. ROCE allows distinguishing between the two strategies by using a temporal, yet spatially homogeneous virtual stimulus. A temporal comparison will elicit a response, but not a spatial one. In the future, ROCE can be applied to chemotactic cell types with unknown sensing mechanism(s). Using the respective cellular messengers, other sensory taxes such as thermotaxis, magnetotaxis, and phototaxis can be studied as well.

ROCE is powerful to generate naturalistic, quantitative temporal stimuli. Owing to their periodic movement and rapid cellular signaling, sperm provide an ideal test system. We gained several insights into sperm chemotaxis. First, oscillating *s*_v_(*t*) induces oscillating voltage responses. Because the sperm CNGK channel that generates the electrical signal does not inactivate or desensitize at high cGMP concentrations (Bönigk et al., 2009), this result indirectly implies that periodic stimulation also elicits phase-locked oscillations of cGMP concentration with a frequency of *s*(*t*). We anticipate that oscillations arise from a balance between cGMP synthesis and hydrolysis. Second, sperm sample a stimulus continuously in time (rather than by intermittent ‘sniffing’) and react almost instantaneously. Third, the signaling pathway operates as low-pass filter that dampens high-frequency fluctuations of *s*(*t*) above 1 Hz. Fluctuations can arise either from small-scale inhomogeneities of chemoattractant distribution, active fluctuations of flagellar swimming (Ma et al., 2014), or shot-noise of chemosensation at low chemoattractant concentrations (Kromer et al., 2018). Fourth, the swimming paths driven by self-entrained oscillations or virtual stimulus oscillations match when *f*_s_ = *f*_0_. Finally, it has been suggested that sensory systems and modalities as diverse as sperm chemotaxis and algae phototaxis can be achieved by the same principle of sensory-motor coupling, known as helical klinotaxis (Crenshaw, 1996). Here, using ROCE, we hijacked sperm to alter its sensory modality from chemotaxis to phototaxis. Thus, the same signaling backbone is suitable for navigation using different sensory modalities.

ROCE can decipher cellular computations at each stage along a signaling pathway for naturalistic sensory fields. Using light-imposed patterns of intracellular activity, the feedback loop between a stimulus, intracellular signaling, and the path **r**(*t*) of a motile cell can be studied on physiological relevant timescales. Such experiments hold promise to understand principles of information self-structuring in cells and microorganisms, implement remote control, and inspire biomimetic navigation strategies for micro-robots.

## ACKNOWLEDGEMENTS

We thank Heike Krause for preparing the manuscript. Financial support by the Deutsche Forschungsgemeinschaft (DFG) via the priority program SPP 1726 “Microswimmers” (U.B.K., L.A., and B.M.F.) and the grants [313904155/SA1785/7-1, 380524518/SA1785/9-1, 214362475/GRK1873/2] (P.S.) are gratefully acknowledged. A.M. was supported by a grant from the Italian Ministry of Education, University, and Research 2010599KRB. We thank Marco Gigante (SISSA Mechatronics Lab) for technical assistance in olfactory neuron experiments.

## AUTHOR CONTRIBUTIONS

L.A., H.H., U.B.K., and R.S. designed the project; H.H., M.B., D.F., U.B.K., A.R., and R.S. performed the stopped-flow experiments; A.R. synthesized the caged compounds; L.A., A.G., F.L., and R.P. performed the motility experiments; L.A., B.M.F., and A.G. developed the theory for sperm motility; A.H.C., A.M., and S.P. designed and performed the electrophysiological experiments on olfactory neurons; V.D., B.K.F., and P.S. designed and performed the experiments on cardiac myocytes. H.H. and H.G.K. designed and performed the experiments on CHO cells. L.A., B.M.F., H.H., and U.B.K. wrote the manuscript; all authors edited and revised the manuscript.

## Supplementary Figures

**Supplementary Figure 1.**
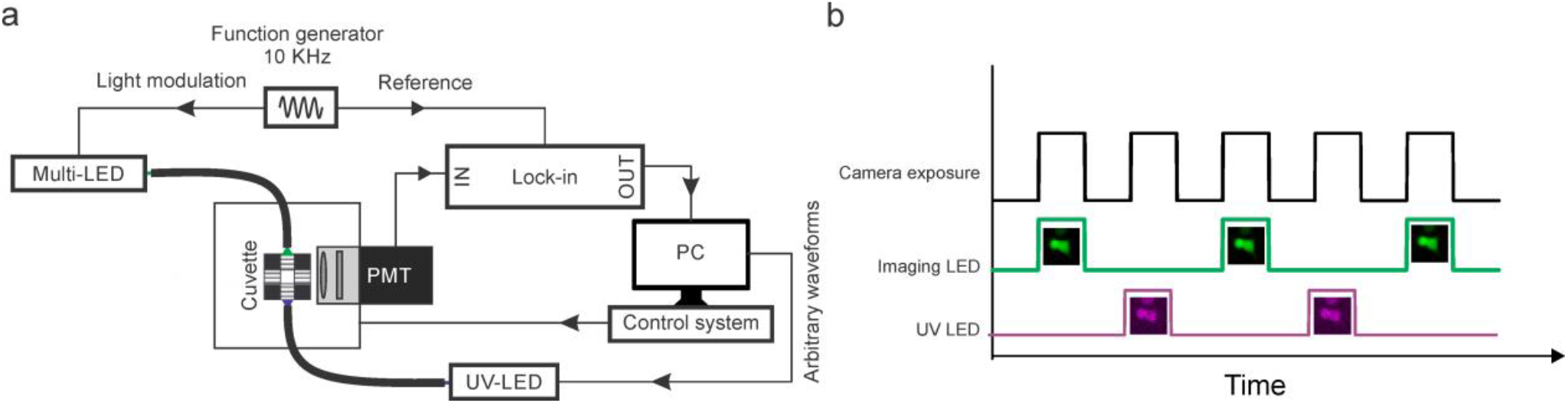
Stopped-flow and imaging setups. **a**, Stopped-flow device operated with the locked-in principle (see methods for detail). PMT refers to a photomultiplier tube. This detection method allowed the simultaneous photo-release of caged compounds and recording of fluorescence. Only the fluorescence signal that was modulated at 10 kHz was recovered because the fluorescence resulting from uncaging was filtered out by the lock-in amplifier. **b**, Imaging protocol used to image cADDis in live cells. Waveforms show the TTL signal used to drive the imaging LED (green) and UV LED (violet) sequentially. This sequential mode allows imaging of fluorescence without interference from photo-release of caged compounds. Odd camera frames were used for quantification of cADDis signal and even frames for the quantification of the release.

**Supplementary Figure 2.**
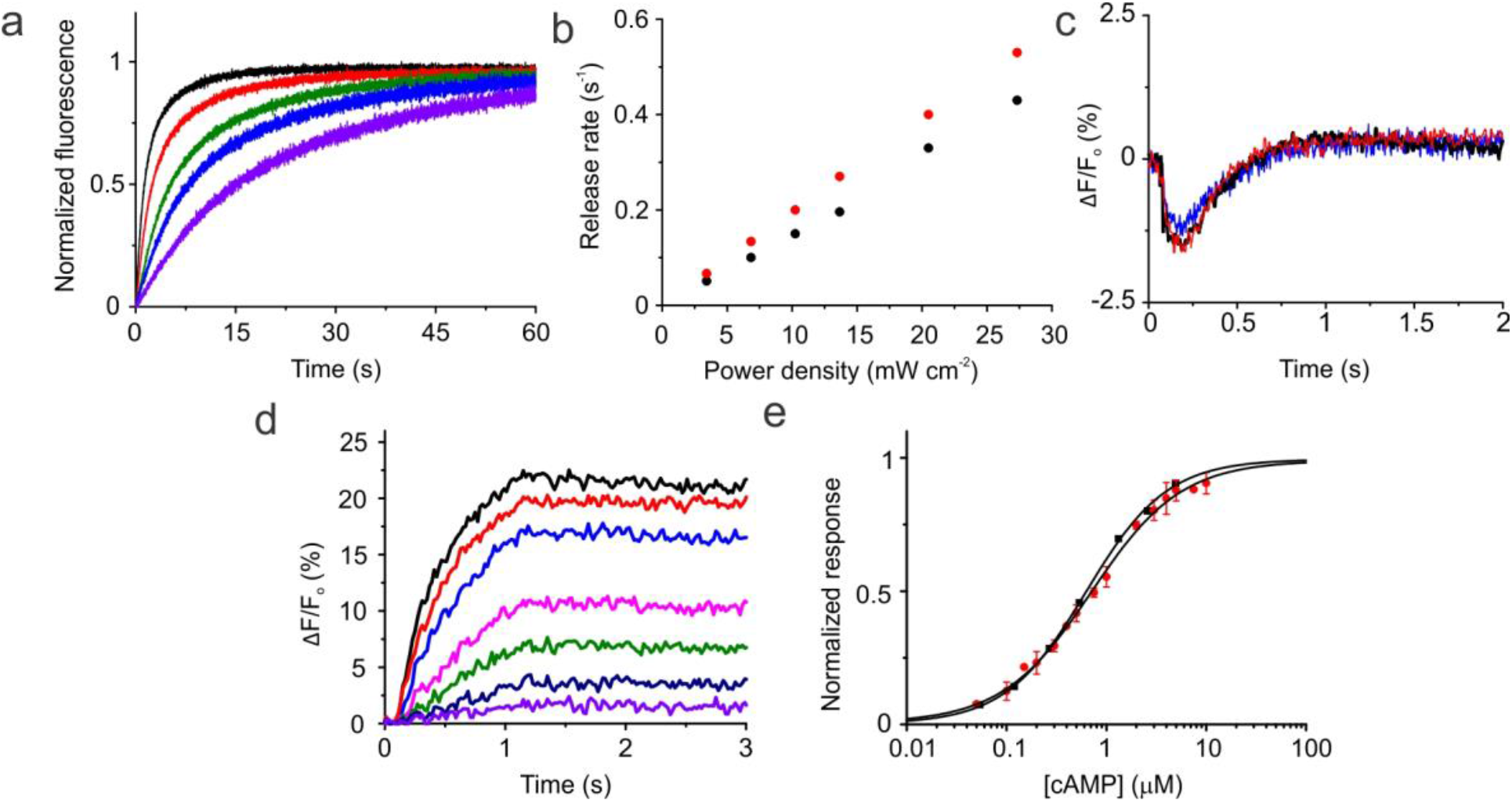
Calibration of photo-release of cNMP. **a**, Time course of DEACM-OH fluorescence during photolysis obtained at different power densities ranging from 3.4 (violet) to 34.2 mW cm^-2^ (black) at 365 nm. **b**, release rate as a function of power density obtained from exponential fit (black) of the fluorescence time courses; theoretical estimate of release rate (red). **c**, Caged cGMP-loading control in sea urchin sperm. Changes in voltage evoked by a 20-nM cGMP release in sperm loaded with different concentrations of DEACM-caged cGMP, 1 μM (black), 15 μM (red), and 25 μM (blue). **d**, Changes in cADDis fluorescence in CHO cells upon photo-release of cAMP: 40 nM (violet), 100 nM (dark blue), 200 nM (green), 400 nM (magenta), 1 μM (blue), 2 μM (red), and 4 μM (black). **e**, Normalized dose-response relation of cAMP to cADDis obtained with two different methods; black dots (stopped-flow recordings from the photo-release of cAMP in live CHO cells), red dots (cADDis in cell lysate, see methods). Fit of a simple binding isotherm to the data (black line) yielded *K*_D_ = 628.3 nM (*n* = 1 experiment) and *K*_D_ = 723.3 ± 45.1 nM (*n* = 3).

**Supplementary Figure 3.**
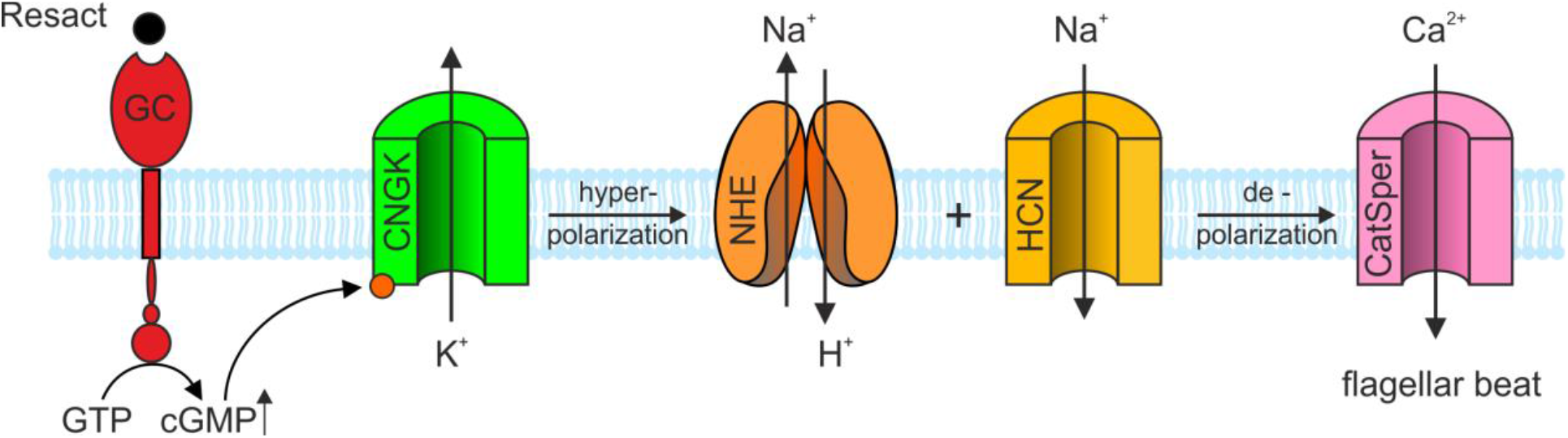
Chemotactic signaling pathway of *A. punctulata* sperm. The icons represent the molecules that encode a chemotactic cell response. GC, guanylate cyclase that synthesizes cGMP and binds the chemoattractant; CNGK, K^+^-selective cGMP-gated ion channel that causes a hyperpolarization; sNHE, Na^+^/H^+^ exchanger gated by hyperpolarization; HCN, hyperpolarization-activated and cGMP-gated pacemaker channel, which carries a depolarizing inward current; CatSper, Ca^2+^ channel gated by changes in pH_i_ and voltage.

**Supplementary Figure 4:**
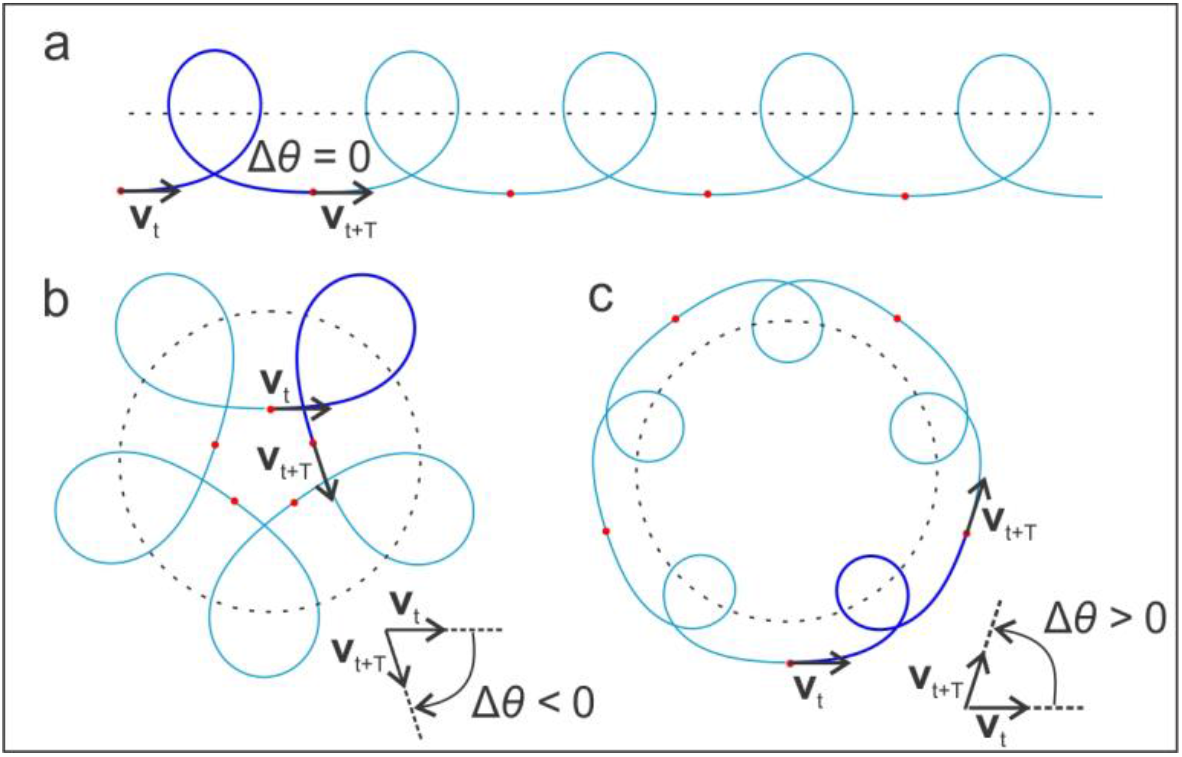
The orientation delay Δ*θ* determines the shape of the swimming path. Dark and light blue lines correspond to the predicted trajectories during the first and the following four stimulus periods, respectively. Red dots indicate the cell position at time points of *t* = *nT*, where *T* is the stimulus period and *n* = 0, 1, 2, 3, 4. Black dash lines intersecting the swimming paths are the centerline of the corresponding path. **v**_t_ is the velocity at time *t*. Each stimulus oscillation results in a curvature oscillation, which in turn produces a lobe shape. **a**, When Δ*θ* = 0, the path is linearly drifting. **b**, Exemplary swimming path for Δ*θ* < 0. **c**, Exemplary path for Δ*θ* > 0. Negative and positive Δ*θ* result in lobes pointing outside or inside of the circle described by the path centerline, respectively.

**Supplementary Table 1.**
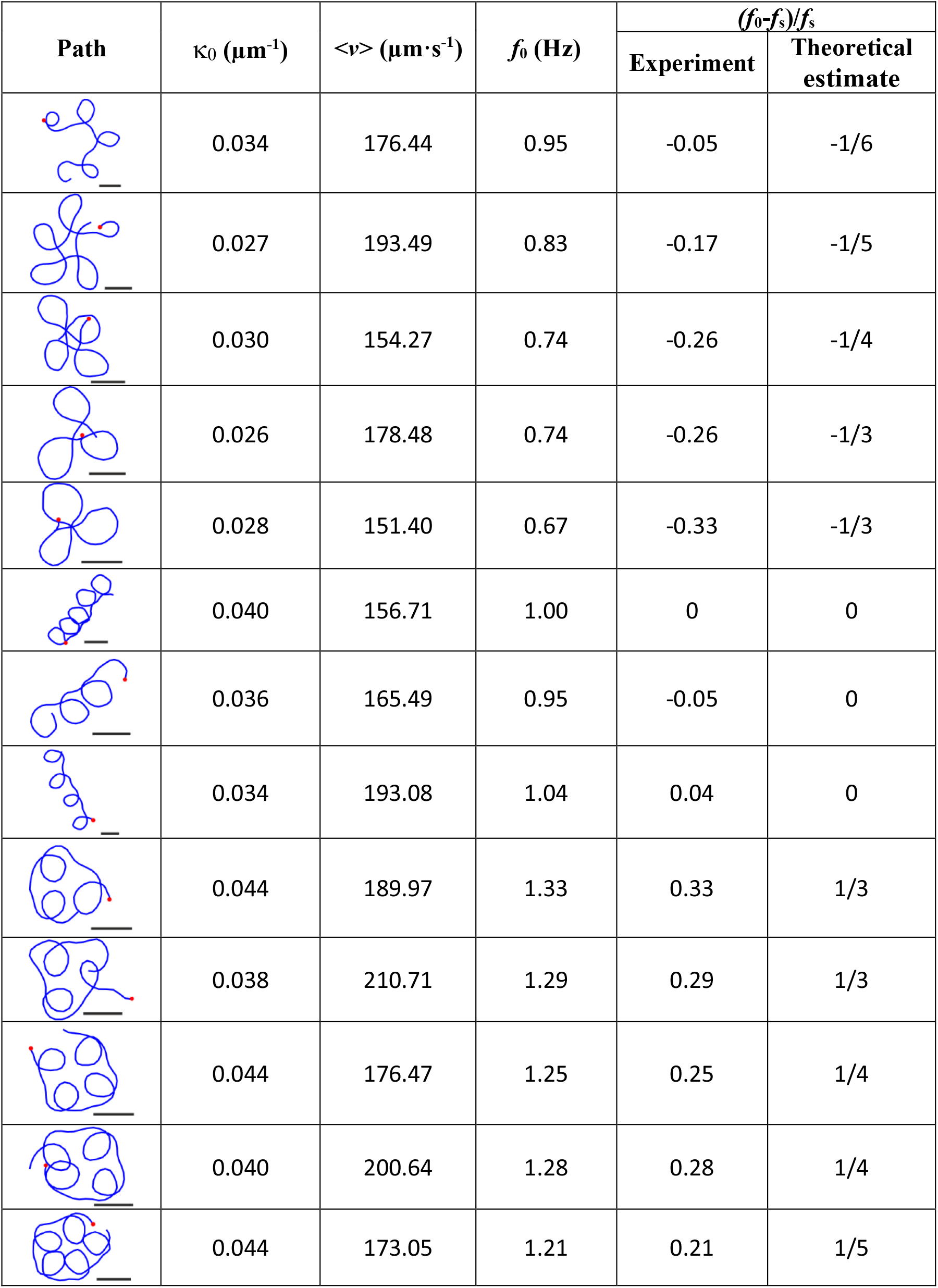
Collection of representative hypotrochoid swimming paths and their corresponding swimming parameters. Swimming paths of cGMP-loaded sperm evoked by a 1-Hz sinusoidal virtual stimulus. Scale bars represent 50 μm. The red dot indicates the starting cell position. The curvature parameter κ_0_ was estimated based on eqn. S4 as κ_0_ = <κ(*t*)>. The intrinsic swimming frequency *f*_0_ is estimated as *f*_0_ = κ_0_·<*v*(*t*)>/2π, where *<v*(*t*)> is the average swimming velocity. Of note, theoretical estimates of *f*_0_ assumed *v* = *v*_0_ constant (*f*_0_ = κ_0_*v*_0_/2π). The associated theoretical ratio (*f*_0_-*f*_s_)/*f*_s_ corresponds to that from a path featuring an integer number of lobes.

| Path | $\kappa_0$ ($\mu\text{m}^{-1}$) | $\langle v \rangle$ ($\mu\text{m}\cdot\text{s}^{-1}$) | $f_0$ (Hz) | $(f_0 - f_s)/f_s$ | |
| --- | --- | --- | --- | --- | --- |
|  |  |  |  | Experiment | Theoretical estimate |
|  | 0.034 | 176.44 | 0.95 | -0.05 | -1/6 |
|  | 0.027 | 193.49 | 0.83 | -0.17 | -1/5 |
|  | 0.030 | 154.27 | 0.74 | -0.26 | -1/4 |
|  | 0.026 | 178.48 | 0.74 | -0.26 | -1/3 |
|  | 0.028 | 151.40 | 0.67 | -0.33 | -1/3 |
|  | 0.040 | 156.71 | 1.00 | 0 | 0 |
|  | 0.036 | 165.49 | 0.95 | -0.05 | 0 |
|  | 0.034 | 193.08 | 1.04 | 0.04 | 0 |
|  | 0.044 | 189.97 | 1.33 | 0.33 | 1/3 |
|  | 0.038 | 210.71 | 1.29 | 0.29 | 1/3 |
|  | 0.044 | 176.47 | 1.25 | 0.25 | 1/4 |
|  | 0.040 | 200.64 | 1.28 | 0.28 | 1/4 |
|  | 0.044 | 173.05 | 1.21 | 0.21 | 1/5 |

**Supplementary Movie 1**. Sea urchin sperm stimulated with a 1 Hz sinusoidal virtual stimulus. Upon stimulation, sperm swim along hypotrochoid paths. The swimming path is displayed before (blue) and during (orange) stimulation. The recording was done using dark-field microscopy and 10x magnification. The virtual stimulus of cGMP release rate is shown. The movie is played at half the real speed.

**Supplementary Movie 2**. Changing the sensory modality of sperm from chemotaxis to phototaxis. Remote control of sea urchin sperm by ROCE in the absence of attractants. Sperm loaded with BECMCM-cGMP are exposed to a Gaussian UV profile for uncaging. The recording was done using dark-field microscopy and 10x magnification. The movie is played at real speed.

**Supplementary Movie 3**. Sperm painting. Sea urchin sperm loaded with BECMCM-cGMP are remote controlled to form a Minerva icon. Upon illumination with a UV light pattern representing Minerva, sperm accumulate in the irradiated area. To increase the resolution, the movie has been made from four recordings containing a quarter of the Minerva icon and have been joined together after the acquisition. Recordings were done using dark-field microscopy and 10x magnification. The movie is played at real speed.

**Supplementary Note 1**. In the following, we calculate the swimming path of sea urchin sperm stimulated by a sinusoidal virtual stimulus releasing cGMP.

### Calculation of path curvature for a sinusoidal stimulus

The sinusoidal UV light stimulus can be described as follows:

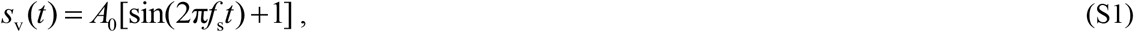

where *f*_s_ is the UV light frequency and *A*_0_ is the stimulus amplitude. This oscillatory UV stimulus will release cGMP directly within the sperm cell and will cascade further signaling events. Sperm cells can faithfully translate the oscillatory UV stimulus to phase-shifted oscillations of intercellular Ca^2+^ concentration ([Ca^2+^]_i_) at the same frequency of stimulation (*f*_s_; Fig. 2e). Intracellular Ca^2+^ is reported by a Ca^2+^ indicator whose relative fluorescence changes (*F*_r_ = Δ*F*/*F*_0_) can be described by:

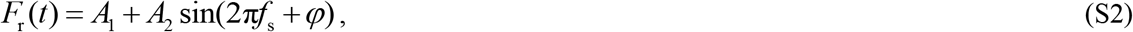

where *A*_1_, *A*_2,_ and ***φ*** are constants and ***φ*** is the phase shift between *s*_v_(*t*) and the oscillatory Ca^2+^ fluorescence signal. The relationship between the swimming path curvature *κ*(*t*) and the Ca^2+^ fluorescence signal *F*_r_ can be described as follows (Alvarez, J. Cell Biology 2012):

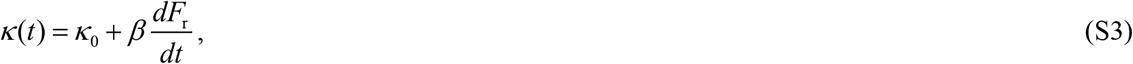

where *κ*_0_ and *β* are constants. By substituting *F*_r_ from equation S2 into S3, *κ* can be calculated as:

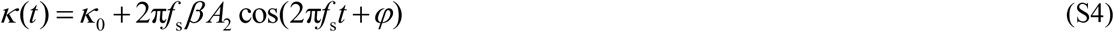

Eqn. S4 states that the path curvature *κ*(*t*) oscillates at the stimulus frequency. Oscillations of *κ*(*t*) result in a lobe shape per stimulus period (Fig. 1).

#### Characterization of the path symmetry

We use the angle *θ* between the velocity vector **v**(*t*) and the abscissa axis to characterize the swimming direction. This angle can be calculated from the geometric definition of curvature *κ*:

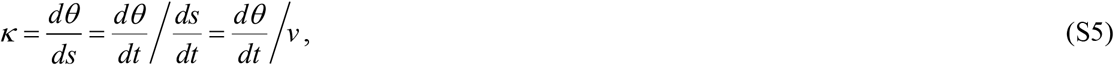

where *v*(*t*)= |**v**(*t*)| is the swimming speed. By inserting *κ* from equation S4 into S5:

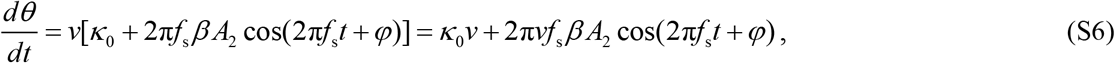

for *v* = *v*_0_ constant, *θ* can be easily calculated:

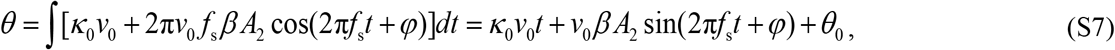

where *θ*_0_ is an integration constant. For a better categorization of the path shape, we introduce a variable named “orientation delay” Δ*θ*, which is the difference between the velocity orientation angle at time *t* and the same angle one stimulus period later (here *T* = 1/ *f*_s_; see Fig. 1):

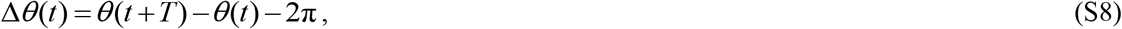

where the arbitrary angular constant 2π is introduced for convenience and takes into account that the expected rotation angle for one stimulus period is about 2π. Using Eqn. S7, Δ*θ* can be calculated:

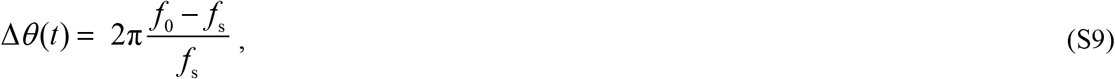

with *f*_0_ = *κ*_0_*ν*_0_/2π the angular rotation frequency of the velocity vector. For integer *N* = 2π/Δ*θ*, the path has a *N* fold-symmetry with *N* = *f*_s_/(*f*_0_-*f*_s_) lobes. For negative Δ*θ*, the path lobes will stretch out of the circle described by the centerline. For Δ*θ* = 0, the paths drift along a line, and for positive Δ*θ*, the path lobes will bend inward of the circle described by path centerline (Supplementary Fig. 4). Finally, the phase delay ***φ*** between the Ca^2+^ signal and the path curvature will affect the orientation of the path without changing the overall shape. This result is consistent with equation S9, which tells us Δ*θ* is only dominated by the ratio of (*f*_0_-*f*_s_)/*f*_s_.

## Methods

### Stopped-flow measurements using the lock-in detection principle

We measured changes in [Ca^2+^]_i_, and voltage V_m_ in a rapid-mixing device (SFM-400; BioLogic) in the stopped-flow mode at 18°C (Hamzeh et al., 2019) (Supplementary Fig. 1). The changes in [Ca^2+^]_i_, and V_m_ were measured with the Ca^2+^ indicator Fluo-4-AM, the pH indicator pHrodo Red, and the voltage-sensitive indicator FluoVolt (Molecular Probes), respectively. Dry sperm were suspended 1:6 (v/v) in loading buffer containing ASW, the respective indicator (10 μM) and 0.5% Pluronic F127 (Sigma-Aldrich or Molecular Probes); loading time was 30–60 min (Fluo-4-AM) or 15-30 min (FluoVolt) at 18°C (for details see (Hamzeh et al., 2019)). After incubation, the sample was diluted 1:20 with ASW containing DEACM-caged cGMP (15 μM, if not otherwise indicated) and 0.25% (w/v) Pluronic F127. The caged cGMP was allowed to equilibrate across membranes for 7 min. The completion of equilibration was ascertained by light flashes that photo-released cGMP and evoked a V_m_ or Ca^2+^ response. After 7 min, the amplitude of V_m_ responses leveled off and remained constant for up to 20 min. Sperm samples were kept in caged cGMP/ASW solution throughout an experiment (about 15 min) to prevent caged compounds leaking slowly from sperm after 1:20 dilution. In the stopped-flow device, the sperm suspension (about 3x10^8^ cells/ml) was rapidly mixed 1:1 (v/v) at a flow rate of 1 ml/s with ASW. Leakage of caged cGMP after 1:1 mixing was negligible during recording times of up to 30 s.

Fluorescence of indicators was excited by a SpectraX Light Engine (Lumencor). The excitation light was modulated at a reference frequency of 10 kHz and directed to the cuvette using a liquid light guide. Emission was recorded by photomultiplier modules (H9656-20; Hamamatsu Photonics) and the signal was recovered using digital lock-in amplifiers (Model 7230, Signal Recovery, Oakridge, TN, USA). The lock-in amplifiers can enhance the signal-to-noise at a given reference frequency while excluding noise and signals at different frequencies. This detection method allowed the simultaneous photo-release of the caged compounds and recording of fluorescence. Only the fluorescence signal that was modulated at 10 kHz was recovered and the fluorescence resulting from uncaging was filtered out. Data acquisition was performed with a data acquisition pad (PCI-6221; National Instruments) and the software Bio-Kine (BioLogic). Fluo-4 fluorescence was excited with a cyan LED with a band-pass filter (ET 490/20; Chroma), and emission was recorded after a band-pass filter (BrightLine 536/40; Semrock). FluoVolt fluorescence was excited with a teal LED with a band-pass filter (BrightLine 513/17; Semrock), and emission was collected after a band-pass filter (BrightLine 542/20; Semrock) for FluoVolt. Signals represent the average of at least two recordings and are depicted as the percent change in fluorescence (Δ*F*) with respect to the mean of the first 5–10 data points before the onset of the signal (*F*_o_). The control (ASW) Δ*F*/*F*_o_ signal was subtracted from the resact-or cGMP-induced signals.

### Experimental determination of photo-release

The experimental estimation relied on the fact that the DEACM cage is only weakly fluorescent when bound to cNMP, but becomes highly fluorescent upon photolysis (Supplementary Fig. 2a) (Eckhardt et al., 2002). The increase in fluorescence was measured upon UV excitation at increasing light levels and the fluorescence curves were normalized by the corresponding light level. The change in fluorescence over time was fitted with an exponential function (Supplementary Fig. 2a) of the form:

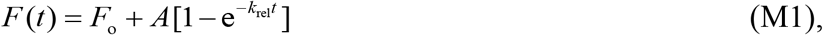

where *F*(*t*) represents the fluorescence at time *t, F*_o_ represents background fluorescence, *A* is a constant, and *k*_rel_ is the release rate. The plot of the slope as a function of power density gave a linear fit that was in line with the theoretical estimation (Supplementary Fig. 2b) and was used to predict the photo-release for various waveforms.

### Theoretical estimate of photo-release

In the following, we derive analytically the release efficiency of caged compounds. Using the Beer-Lambert law, the attenuation of UV light while passing through a solution containing a caged compound at a concentration *c* is:

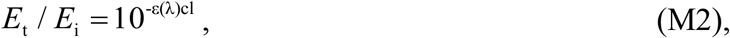

wherein *ε*(*λ*) is the extinction coefficient and *l* is the travelling path length of light in the solution; *E*_i_ and *E*_t_ refer to the incident and transmitted light energy, respectively. For a typical *ε*(*λ*) of 10,000 M^-1^cm^-1^ (Goeldner and Givens, 2005), a compound concentration of *c* ∼ 10 μM, and a path length *l* ∼ 0.1 cm, the exponential factor *ε*(*λ*)*cl* is ∼ 0.01 << 1, and eqn. M2 can be linearized by Taylor expansion to:

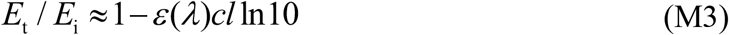

Using the Planck-Einstein relation and eqn. M3, the number of absorbed photons *n*_a_ for a given wavelength *λ* can be estimated:

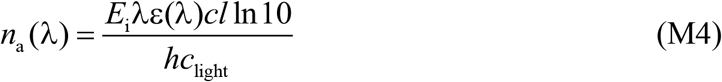

wherein *c*_light_ is the speed of light and *h* is the Planck constant. The fraction of absorbed photons that trigger a photochemical reaction *n*_r_ is given by ϕ*n*_a_, wherein ϕ is the quantum yield of photolysis. Thus, *n*_r_ is equal to the number of released molecules. Considering the spectral flux of the light source *P*(λ), and the flash duration *t*. For a narrow spectral band d*λ, n*_r_ is:

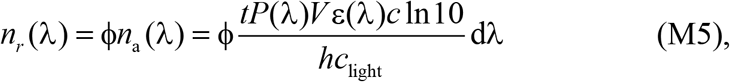

where *V* is the irradiated volume. The number of caged molecules before the flash in that volume is *n*_*o*_ = *VcN*_A_ with *N*_A_ being the Avogadro number. Integrating for all wavelengths, the fraction of released molecules per unit of time is:

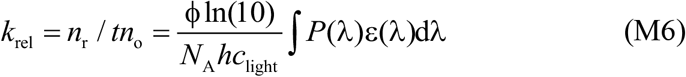

Assuming that the release is a first order reaction of the caged compound with a photon of the form:

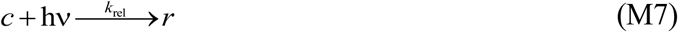

with released compound concentration *r*. The mass balance equation, and the principle of mass conservation stay that:

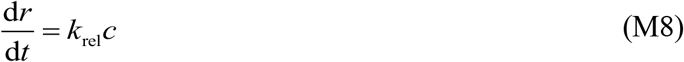

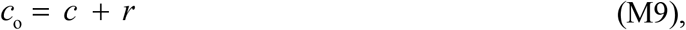

where *c*_o_ is the caged compound concentration before photolysis. The solution for this equation is, assuming *k* constant:

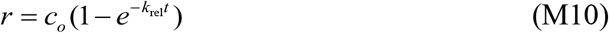

Combining eqn. M6 and M10, the total release can be quantified:

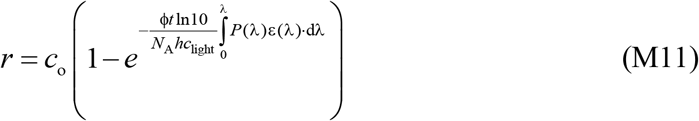

If the light spectral flux varies with time (*P* = *P*(*t*,λ)), a more general solution to eqn. M8 has to be used instead, and the following expression can be readily found:

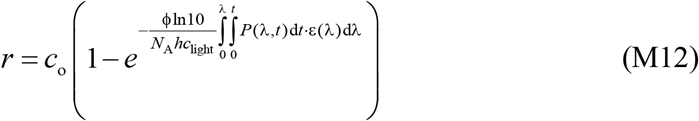

A graphical user interface written in MATLAB and LabVIEW to evaluate the release of several compounds using different light sources and illumination protocols can be found here.

### Calibration of intracellular caged cNMP concentrations and fractional cNMP release

To calibrate ROCE for precise quantification of cell signaling, the intracellular caged-cNMP concentrations and fractional photolysis were determined experimentally and theoretically (Supplementary Fig. 2). Theoretical and experimental estimates agree within 30% (Supplementary Fig. 2b).

DEACM-caged cNMPs are neutral and readily cross cell membranes; unlike acetomethoxy esters, they should not accumulate inside cells. Consequently, extra- and intracellular cNMP concentrations should be approximately equal. We tested this premise in two different cell systems using two different methods. First, sperm were loaded with 1-25 μM DEACM-caged cGMP. The photolytic cGMP release was kept constant by adjusting the light energy accordingly. Sperm respond to stimulation by chemoattractant or cGMP with a hyperpolarization of membrane potential V_m_ (Strünker et al., 2006). The cGMP-evoked V_m_ responses perfectly superimposed (Supplementary Fig. 2c), demonstrating that external and cytosolic caged compound concentrations linearly scale or are equal. As a second test, we determined by light titration the *K*_D_ constant of cAMP binding to a cAMP sensor, named *cAMP Difference Detector in situ* (cADDis) (Moore et al., 2016), in CHO cells (Supplementary Fig. 2d, e). The cADDis sensor was calibrated by two different procedures. Viral transduction of the cAMP sensor was performed according to the manufacturer’s protocol (Red Fluorescent cAMP assay, Montana Molecular). In the first procedure, 4 × 10^6^ cells of a stable cell line expressing mlCNBD-FRET, another cAMP biosensor (Mukherjee et al., 2016), were mixed with the cADDis BacMam stock, complete medium, and sodium butyrate and plated on a 9-cm cell culture dish. Cells were incubated for 30 min at room temperature in the dark and then for 48 h at 37°C and 5% CO_2_. Cells were suspended in 500 μl hypotonic medium containing (10 Mm Hepes/NaOH) at pH 7.4 and 2 mM EDTA and sonicated three times for 10 sec to lyse the cells. After centrifugation (500xg, 10 min), the supernatant was resuspended in 5 ml solution containing (in mM): NaCl 140, KCl 5.4, MgCl_2_ 1, CaCl_2_ 1.8, glucose 10, and Hepes 5 adjusted to pH 7.4 with NaOH. We recorded the fluorescence emission spectrum of cADDis (565-650 nm) excited at 550 nm. The cAMP was added stepwise, and spectra were again recorded. As a reference, we recorded from the same sample for each cAMP concentration the emission spectrum (460-580 nm) of mlCNBD-FRET, excited at 430 nm. Fluorescence values were plotted vs. cAMP concentration and analyzed using a simple binding model. The *K*_D_ value for cAMP binding to cADDis and mlCNBD-FRET was 723.3 ± 45.1 nM (*n* = 3) and 144.7 ± 15.0 nM (*n* = 3), respectively. Second, cADDis was calibrated *in situ* by defined photolytic release of cAMP from caged cAMP-loaded CHO cells. The changes in cADDis fluorescence were fit with a simple binding model using Origin (OriginLab). The KD values agree well with each other (723.3 nM vs. 628.3 nM). Thus, cNMP photo-release inside cells can be precisely determined within live cells.

### cADDis imaging

HEK cells expressing cADDis and loaded with DEACM-caged cGMP (15 μM), were bathed in ES containing 0.01% Pluronic F127 (Sigma-Aldrich) and imaged in a custom-made observation chamber. Cyclic cAMP was released by a 365 nm UV LED (M365L2; Thorlabs) with a custom-made power supply whose current was set by a data acquisition card (PCI-6040E; National Instruments). The output power was adjusted in time using a customized LabVIEW program. The UV light was coupled into the back port of an inverted microscope (IX 71; Olympus) equipped with a 10x objective and with a 530LP dichroic (T525 LPXR; Chroma) and a band pass emission filter (BrightLine 562/40; Semrock). An EMCCD camera (iXon EM + DU-897; Andor Technology) was used to collect fluorescent images. Sequential imaging was used in order to separate cADDis fluorescence from the background fluorescence of the released cage into sequential camera frames (Supplementary Fig. 1b).

### Data analysis of cADDis fluorescence

The concentration of total cAMP released at any time *t* can be represented as:

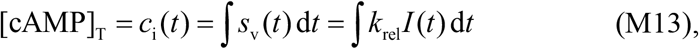

where *s*_v_(*t*) represents the virtual stimulus function, *k*_rel_ the photorelease rate, and *I*(*t*) the light intensity. The cADDis time traces were fit with a modified simple binding isotherm using Origin (OriginLab) of the form:

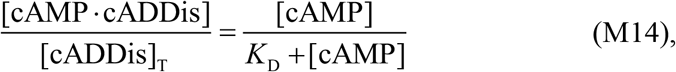

where [cADDis]_T_ is the total cADDis concentration, [cAMP] is the free cAMP concentration, and [cAMP·cADDis] is the concentration of the complex between cAMP and cADDis. Assuming that the fluoresence signal is proportional to the complex concentration, then the change in fluorescence of cADDis can be described as:

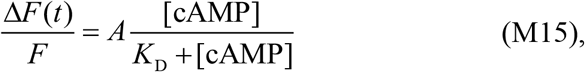

where *A* is a proportionality factor. Finally, if only a small fraction of released cAMP binds to the sensor cADDis, free and total cAMP are approximately equal, and eqn M13 and M14 can be combined to give:

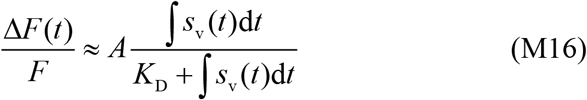

The cADDis fluorescence time course was fitted to the above equation for different stimulus functions: *s*_v_(*t*) = *s*_1_, *s*_v_(*t*) = *s*_2_*t*, and *s*_v_(*t*) = *s*_2_sin(ω*t*) + *s*_1_, corresponding to a step, a ramp, and a sinusoidal waveform, respectively.

### Single-cell recordings using temporal UV-light waveforms

Sperm loaded with DEACM-caged cGMP (15 μM), were diluted 1:10^6^ in ASW containing 0.5% Pluronic F127 (Sigma-Aldrich) and imaged in a custom-made observation chambers of 100-μm depth. Cyclic GMP was released by a 365 nm UV LED (M365L2; Thorlabs) with a custom-made power supply whose current was set by a data acquisition card (PCI-6040E; National Instruments). The output power was adjusted in time using a customized LabVIEW program. The UV power was calibrated using a photodiode (DET36A/m; Thorlabs). The UV light was coupled into the back port of an inverted microscope (IX 71; Olympus) equipped with a 10x objective (UPLSAPO10X; Olympus) and with a 495LP dichroic (BS495 LP; Semrock). UV light was attenuated by a 2% neutral-density filter (Linos, Qioptiq Photonics) and filtered for imaging by a 510 LP emission filter (ET510LP; Chroma). The maximal light power delivered onto the sample (20 μW) was measured with a power meter (Fieldmax top and PowerMax PS19Q; Coherent). Dark-field illumination was achieved with a 730-nm LED (M730L4; Thorlabs) and a short working distance condenser (IX2-U-UCD8; Olympus). Dark-field images were recorded by an EMCCD camera (iXon EM+ DU-897; Andor Technology).

### Transfer function calculation

The frequency sweep input was generated using the MATLAB chirp function, with a instantaneous frequency sweep given by:

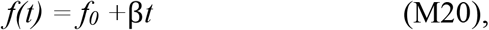

where: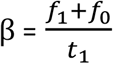, with *f*_0_ is the instantaneous frequency at time 0, and *f*_1_ is the instantaneous frequency at time *t*_1_.

Multiple chirp *s*_v_(*t*) inputs were used sequentially on sperm from the same animal in order to cover a range between 0 Hz and 15 Hz with an overlap in the frequency ranges of the inputs, the multiple inputs produced multiple outputs that were later used to extract a set of impulse responses using convolution theorem:

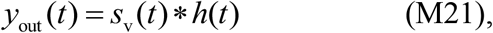

where *y*_out_ represents the measured Ca^2+^ output, ∗represents the convolution, and *h* the transfer function. The convolution theorem states that the Fourier transform of a convolution of two signals is the pointwise product of their Fourier transforms, therefore equation M21 could be written as:

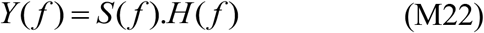

*Y* is the Fourier transform of *y*_out_, *S* is the Fourier transform of *s*_v_, *H* is the Fourier transform of *h*, and *f* is the frequency. The magnitude and phase of the transfer function *H* were calculated as follows:

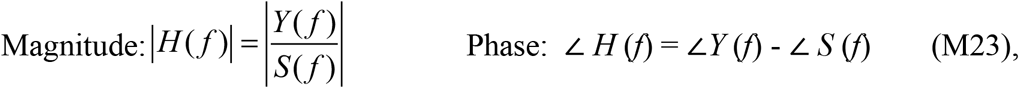

where ∠*H*(*f*), ∠*Y*(*f*), and ∠*S*(*f*) are the phase spectrum of the transfer function, the output, and the input, respectively. The magnitude of the transfer function can be converted into the dB scale by *H* _dB_ (*f*) = 20 log_10_ |*H* (*f*) | . The MATLAB output was in radians. Finally, the set of transfer functions from the multiple chirps were averaged.

The transfer function was tested by providing a sinusoidal stimulus input carrying two frequencies (1 and 6 Hz), and the Ca^2+^ output was predicted as follows:

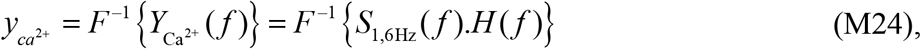

where *F*^-1^ is the inverse Fourier transform. The predicted output was in agreement with the measured Ca^2+^ signal (Fig 3d). All calculations where done using MATLAB single processing toolbox in a custom-made GUI. Data was post-processed and plotted using OriginLab.

### Analysis of spontaneous beating of cardiomyocytes

Neonatal cardiomyocytes were harvested from P2-P4 CD1 mice using the neonatal heart dissociation kit, mouse and rat according to manufacturer’s instructions (MACS Miltenyi Biotec, but without the gentleMACS Dissociator). Dissociated cells were resuspended in Iscove’s Modified Dulbecco’s Medium (IMDM, Thermofisher) supplemented with 20% FCS (PAN-Biotech), 0.1 mM nonessential amino acids, 100 U ml^-1^ penicillin, 100 mg ml^-1^ streptomycin, 2 mg ml^-1^ L-glutamine (all Thermofisher) and plated at low densities on 0.1% gelatin coated coverslips. After 24 h medium was replaced by IMDM medium with 2% FCS. Frequency measurements were performed 3-7 days after cell isolation in Tyrode solution (in mM): 142 NaCl, 5.4 KCl, 1.8 CaCl_2_, 2 MgCl_2_, 10 glucose and 10 Hepes; pH 7.4) containing 20 μM DEACM-caged cAMP and 0.1% Pluronic F127 at 37°C. Spontaneous beating of single neonatal cardiomyocytes was imaged using infrared LED (760 nm) and a CCD camera (piA640-210gm, Basler). Beating frequency was analyzed online using custom designed software (LabVIEW, National Instruments) as reported earlier (Bruegmann et al., 2010). Data was recorded and analyzed with a Powerlab system and the LabChart software (ADInstruments).

DEACM uncaging was performed by illumination through the objective (10x Fluar, Zeiss) with a 365-nm LED (Omicron LEDMOD LAB) operated by a signal generator (Model 2100, A-M Systems) and coupled to an Axiovert 200 microscope (Zeiss) with an optical fiber. Light intensity was calibrated at the level of cells with a power meter (PM100A power meter and S170C sensor, Thorlabs) and adjusted to 6.3 mW mm^-2^. Cardiomyocytes were stimulated every 60 sec with increasing illumination times (1 -250 ms). Baseline and response frequency was determined as mean frequency in a three sec interval two sec before and after illumination, respectively. Cells that did not show a frequency increase and cells with large frequency fluctuations during baseline or between stimulations were excluded from the statistical analysis.

Magnitude of cAMP release by each light pulse was calculated as described above. Dose (log cAMP concentration) versus response (beating frequency) curve was calculated for each individual cell isolation (*n* = 3) and fitted with a nonlinear regression (Graphpad Prism).

### Electrophysiological recordings of olfactory sensory neurons from acute slices of the mouse olfactory epithelium

Coronal slices were prepared from olfactory epithelia of mouse pups (P0–P4) as previously described (Pietra et al., 2016). Mice were decapitated and the nose was dissected en bloc and embedded in 3% low-grade agar (A7002, Sigma) prepared with Ringer’s saline solution containing (in mM): 140 NaCl, 5 KCl, 2 CaCl_2_, 1 MgCl_2_, 10 Glucose and 10 Hepes adjusted to pH 7.4 with NaOH. A vibratome was used to cut coronal slices of 300-μm thickness. During the cutting, the tissue was completely submerged in Ringer’s solution continuously oxygenated and kept near 0°C. After sectioning, coronal slices were allowed to recover for at least 30 min in the same solution.

Currents were measured with a Multiclamp 700B patch-clamp amplifier (Molecular Devices) in the whole-cell voltage clamp mode. Patch pipettes were made using borosilicate capillaries (WPI) and pulled with a Narishige PP10 puller (Narishige). The pipette resistance was 5–7 MΩ when filled with the intracellular solution containing in (mM): 80 K-gluconate, 60 KCl, 2 Mg-ATP, 0.5 EGTA, 10 Hepes, and 0.02 DEACM-cAMP. Currents were low-pass filtered at 2 kHz and acquired at 10 kHz by the analogue-to-digital interface Digidata 1440 (Molecular Devices). All experiments were performed at room temperature (20°–22°C). Cyclic AMP was photoreleased by a 405 nm UV LED (M405L3; Thorlabs) with a LED driver (LEDD1B; Thorlabs) controlled by Clampex 10 (Molecular Devices) through the interface Digidata 1440. The UV light was coupled with a custom-made multi-LED illumination system into the epifluorescence path of an upright microscope (IX51WI; Olympus) equipped with a 60X objective. UV light was first focused to the image plane with a 500-μm pinhole (P500H; Thorlabs). The UV light was reflected initially by a dichroic mirror (T425lpxr; Chroma) and then with a multiband dichroic mirror (69010bs; Chroma). The illumination spot was about 9 μm in diameter in the specimen focal plane. Alexa Fluor 594 (A33082; Thermo Fisher Scientific) was added to the pipette solution to a final concentration of 10 μg ml^-1^ to visualize and locate the cilia and knob of the olfactory neurons. The samples were illuminated with a 565-nm LED (M565L3; Thorlabs) coupled to the custom-made multi-LED illumination system. The 565-nm light was filtered by a 560-nm excitation filter (ET560/25x; Chroma) and reflected by the multiband dichroic mirror (69010bs; Chroma). Fluorescence from Alexa Fluor 594 was filtered with a multiband emission filter (69010e; Chroma) and visualized by a CMOS camera (DFK72BUC02; Imaging Source). Both, dendritic knob and cilia of the recording neuron were placed under the UV spot region for cAMP release. The interval between light stimulations was at least 60 s to allow recovery from adaptation (). Light power was measured with a microscope-slide power meter (S170C; Thorlabs).

### Path simulation

All theoretical trajectories were calculated numerically using eqn. S4 and by time integration of the geometric definition of curvature (eqn. S6) (Supplementary Note) with parameters: κ_0_ = 12 mm^-1^, β = 0.04 s mm^-1^ (Alvarez et al., 2012), *f*_s_ = 1 Hz, and *A*_2_ = 40.

### Remote control of single cells

The UV light gradient was generated by a Digital Light Processing (DLP) beamer with a 385 nm wavelength (Pro4500; Wintech Digital) collimated by one eyepiece lens (W10x/23; Carl Zeiss) and focused into the focal plane of the sample by a plano-convex lens (ACA254-150-A *f* = 150 mm; Thorlabs). A 425LP dichroic (425DCXR; Chroma) unde r the objective lens (UPLSAPO10X; Olympus) was used to couple the UV light onto the infinity-corrected light path of the microscope (IX 71; Olympus). For estimating and correcting the UV gradient shape after passing all the optical components, a thin sample solution of sodium fluorescein (200 μM; Fluka Analytical) was imaged. Fluorescence emission was filtered by a 535/30 filter (ET535/30M; Chroma) and used to calculate a height-width ratio deformation factor, which was applied to all images projected by the DLP beamer. For loading, dry sperm were diluted 18:100 (v/v) in ASW containing 30 μM [6,7-bis(ethoxycarbonylmethoxy)-coumarin-4-yl]methyl-cGMP (BECMCM-caged cGMP; (Hagen et al., 2001; Hagen et al., 2002), 2.5 mM sodium Probenecid (AAT Bioquest), 0.5% Pluronic F127 and kept in the dark at 20°C for 1 hour. After loading, sperm were studied at 1:10^5^ dilution in ASW containing 30 μM BECMCM-caged cGMP, 2.5 mM Probenecid, and 0.5% Pluronic in a custom-made observation chamber with a 150-μm depth. Dark-field images were generated with a 660 nm LED light source (M660L3-C1; Thorlabs) and recorded (100 fps) by a fast monochrome camera (Dimax; PCO) after UV-light filtering by a 675/67 filter (FF02-675/67-25; Semrock). To increase the resolution of the Minerva shape formed by sperm, the Minerva stimulus was split into four parts, and the recorded videos were later connected.

